# EEG correlates of physical effort and reward processing during reinforcement learning

**DOI:** 10.1101/2020.01.09.900530

**Authors:** Dimitrios J. Palidis, Paul L. Gribble

## Abstract

Effort-based decision making is often described by choices according to subjective value, a function of reward discounted by effort. We asked whether a neural reinforcement learning signal, the feedback related negativity (FRN), is modulated not only by reward outcomes but also physical effort. We recorded EEG from human participants while they performed a task in which they were required to accurately produce target levels of muscle activation to receive rewards. Participants performed isometric knee extensions while quadriceps muscle activation was recorded using EMG. Real-time feedback indicated muscle activation relative to a target. On a given trial, the target muscle activation required either low or high effort. The effort was determined probabilistically according to a binary choice, such that the responses were associated with 20% and 80% probability of high effort. This contingency could only be known by experience, and it reversed periodically. After each trial binary reinforcement feedback was provided to indicate whether participants were sufficiently accurate in producing the target muscle activity. Participants adaptively avoided effort by switching responses more frequently after choices that resulted in hard effort. Feedback after participants’ choices which revealed the resulting effort requirement for the subsequent knee extension did not elicit an FRN component. However, the neural response to reinforcement feedback after the knee extension was increased during and after the time period of the FRN by preceding physical effort. Thus, retrospective effort modulates reward processing which may underlie paradoxical behavioral findings whereby rewards requiring more effort to obtain can become more powerful reinforcers.

**Significance Statement:** When making decisions, we typically select more rewarding and less effortful options. Neural reinforcement learning signals reinforce rewarding actions and deter punishing actions. When participants received feedback that their choices would require easy or hard physical effort, we did not observe reinforcement learning signals that are typically observed in response to feedback predicting reward and punishment. Thus, the reinforcement learning system does not strictly treat effort as loss or punishment. However, when the effort was completed and participants received feedback indicating whether they successfully achieved a reward or not, reinforcement learning signals were amplified by preceding effort. Thus, retrospective effort can affect neural responses to reinforcement outcomes, which may explain how effort can enhance the motivational effect of reinforcers.

## Introduction

Humans and other animals tend to make decisions that lead to more rewarding and less physically effortful outcomes (Shadmehr et al. 2016; Morel et al. 2017; Walton et al. 2006; Rangel and Hare 2010; Selinger et al. 2015; Kennerley et al. 2009). When learning about reward contingencies in uncertain or changing environments, midbrain dopaminergic neurons signal the difference between expected and obtained reward to the ventral striatum and prefrontal regions (Puig and Miller 2012; Graybiel 2008; Schultz 2006; Holroyd and H 2002; Pessiglione et al. 2006; Bayer and Glimcher 2005; Gläscher et al. 2010). This reward prediction error signal is thought to drive reinforcement learning by updating reward expectations. Dopaminergic, prefrontal and striatal processes have also been implicated in signaling effort cost and motivating effortful behavior to obtain reward (Hosking et al. 2015; Denk et al. 2005; Kurniawan et al. 2011; Kurniawan et al. 2010; Walton et al. 2003; Schweimer et al. 2005; Rudebeck et al. 2008; Salamone et al. 2007; Salamone et al. 2003). However, it remains unclear whether adaptive reward maximization and effort minimization are implemented by an overlapping reinforcement learning circuit or whether they are subserved by distinct but possibly isomorphic neural processes.

During decision making the anterior cingulate cortex (ACC) is known to encode prospective reward and effort cost and to integrate both into a unitary subjective utility signal (Klein-Flügge et al. 2016; Croxson et al. 2009; Kennerley, Behrens, and Wallis 2011; Porter, Hillman, and Bilkey 2019). During outcome evaluation the ACC encodes reward prediction error and supports reinforcement learning (Kennerley, Behrens, and Wallis 2011; Walsh and Anderson 2012a; Amiez, Joseph, and Procyk 2005; Williams et al. 2004; Ito et al. 2003). Thus, the ACC is a possible substrate for a unitary learning signal that integrates reward and effort to shape future behavior. Contrary to this idea, fMRI studies have argued that separate neural systems underlie reward and effort learning, with ACC activity reflecting prediction errors for effort but not reward (Hauser, Eldar, and Dolan 2017; Skvortsova, Palminteri, and Pessiglione 2014). However, an event related potential measured by EEG called the feedback related negativity (FRN), or alternatively the reward positivity, is a reliable neural correlate of reward prediction error and is consistently localized to the ACC (Walsh and Anderson 2012b; Holroyd and H 2002; Holroyd, Krigolson, and Lee 2011; Becker et al. 2014; Vezoli and Procyk 2009). Thus, we sought to test whether the FRN not only acts as a learning signal for reward outcomes but also integrates effort requirements during learning.

Economic theories assert that effort is a cost which devalues reward, and thus predict a diminished neural response to reinforcement for more costly rewards (Shadmehr et al. 2016; Hauser et al. 2017; Botvinick et al. 2009; Hartmann et al. 2013). Paradoxically, it has been found in humans and animals that effort can enhance the reinforcing quality of rewards (Inzlicht, Shenhav, and Olivola 2018; Lydall, Gilmour, and Dwyer 2010; Zentall 2010; Clement et al. 2000). It may be that prospective effort devalues reward, while retrospective effort amplifies reinforcement. In the present study effort interacted retrospectively with reward. Like many real-world situations, uncertain reward was obtained only after effort expenditure.

Participants first made binary choices, then they received feedback about the resulting effort requirements which were probabilistic and uncertain. Subsequently, they performed an effortful EMG production task for which they received variable reward that was dependent on precisely producing a target level of EMG activity. This trial sequence allowed us to test the hypothesis that effort information is integrated or otherwise remembered during the course of an action, and that this information is used retrospectively to compute subjective utility upon completing the action and observing the reward outcome. According to this hypothesis, feedback indicating effort requirements in the current study would not elicit neural reinforcement signals such as the FRN, whereas the neural response to reward feedback at the end of each trial would be modulated both by reinforcement outcome and the preceding effort. Alternatively, if effort is treated simply as an aversive stimulus or an economic loss by a standard temporal difference learning process, then feedback which predicts the upcoming effort but not the reward outcome should elicit neural reinforcement signals (Mulligan and Hajcak 2018).

We found that effort feedback did not elicit an FRN response. However, the differential neural response to reward and non reward outcomes was enhanced by preceding physical effort during and after the time period of the FRN.

## Material and Methods

### Participants

A total of *n* = 18 healthy participants were included in our study (Mean age: 22.12 years, SD: 3.66, 9 females). Four participants underwent the experimental procedure but were excluded due to excessive EEG artefacts caused by sweat or movement associated with the task. Participants provided written informed consent to experimental procedures approved by the Research Ethics Board at The University of Western Ontario.

### Experimental Setup

To allow for isometric contractions of the quadriceps muscles, participants were restrained to a chair by straps on their shoulders and waists. Participants’ ankles were strapped to a rack fixed at the base of the chair, with the knees bent at approximately 90 deg. Participants were seated in front of a CRT monitor with their hands resting on a table positioned to make button presses on a response box.

### EMG and EEG Recording

Unreferenced EEG activity was recorded at 512 Hz using a 64-channel Biosemi ActiveTwo system (Biosemi). Electrodes were mounted in an elastic cap and distributed according to the extended 10-20 system with electrode Cz placed over the vertex. Instead of the typical ground electrode, Biosemi forms a feedback loop between an active Common Mode Sense electrode and a passive Driven Right Leg electrode. The Common Mode Sense electrode was located in the center of the area between P1, Pz, PO3, and POz. The Driven Right Leg electrode was located in the center of the area between Pz, P2, PO3, and PO4. EOG was recorded with electrodes placed above and below each eye and the outer canthus of each eye. Additional electrodes were placed on each mastoid.

EMG activity was recorded at 2400 Hz bilaterally from the rectus lateralis muscles of the quadriceps using an active electrode system and amplifier (g.USBamp; g.tec Medical Engineering). Two electrodes were placed on each muscle belly for bipolar recordings, and a ground electrode was placed on the left shin. EMG signals were filtered at the time of recording using a 5-500 Hz bandpass filter and a 60 Hz notch filter.

### Visual Feedback of EMG

The EMG signal used to provide online visual feedback of quadriceps muscle activity was first rectified, lowpass filtered with a 10 Hz cutoff frequency, and then down-sampled to 120 Hz. At the beginning of each block, participants performed isometric knee extensions with maximum effort continuously for 4 seconds. All samples greater than the median value recorded during maximum effort were averaged to determine the value of maximum voluntary contraction (MVC) used throughout the block. Subsequently, participants were cued to remain completely still and keep their legs relaxed for 4 seconds. The mean EMG signal during this period was used as a baseline value throughout the block.

During each trial, an animation of a thermometer was displayed to participants. The fluid level of the thermometer increased in real time (monitor refresh rate: 60 Hz) as a linear function of the processed EMG signal. In “hard” effort trials, the top of the thermometer corresponded to 85% of MVC, and the bottom of the thermometer corresponded to the baseline measure. In “easy” effort trials, the top of the thermometer corresponded to 15% of MVC. The baseline measure was made to correspond to the point halfway up the thermometer for the “easy” condition in order to reduce the gain of feedback. The fluid level was calculated separately for each leg based on their respective MVC and baseline measures, and the average was used to display feedback. A running average of fluid level for the previous 60 samples was drawn to the screen to provide smooth feedback. During each trial, the maximum fluid level for that trial was continuously displayed such that the fluid level only increased, and if the participant relaxed their quadriceps muscles the feedback would remain at the same level. This allowed for smooth, ballistic isometric contractions. It also made it so that participants were not required to hold the fluid level constant without visual feedback, which often resulted in the fluid level fluctuating or drifting away from the target during pilot experiments.

### Experimental Task

Participants first performed a block of 28 practice trials (see below). Participants then performed 4 blocks of 74 trials with self-paced rest periods between blocks. Each block consisted of 12 control condition trials, followed by 50 experimental condition trials, and finally 12 additional control condition trials. At the end of each block, participants were verbally surveyed as to how physically effortful the easy and hard effort trials were using a scale of 1 to 5, with 5 corresponding to maximum effort.

#### Experimental Condition

During each trial, participants made a binary choice which probabilistically determined whether the trial would require easy or hard physical effort. The effort contingencies had to be learned through experience. Participants then performed isometric knee extensions to control visual EMG feedback on a screen. Participants were instructed to exceed a minimum level of muscle activation indicated by a visual target while remaining as close as possible to the target. Binary reinforcement feedback was provided at the end of each trial to indicate success or failure, which corresponded to a small monetary reward.

Visual stimuli are shown in Figure 1. An animated thermometer was drawn on the screen throughout the task. A cross was drawn at the top of the thermometer to serve as a target for EMG feedback. Letters “A” and “B” drawn to the left and right of the thermometer represented the options for binary choices made in each trial. Participants initiated each trial by pressing either a left or right button on a response box using their left or right index finger, respectively. Immediately each button press, the choice was indicated by a box appearing around the letter “A” or “B” for the left and right response buttons, respectively. The box remained throughout the trial. One second after the button press, either the word “easy” or the word “hard” replaced the target cross for 700 ms to indicate the effort condition of the trial. The effort condition was determined probabilistically by the participants’ response, and the effort contingencies had to be learned through experience. One of the responses led to a “hard” effort trial with a probability of 0.8, and an “easy” effort trial with a probability of 0.2. The other response led to a “hard” effort trial with a probability of 0.2, and an “easy” effort trial with a probability of 0.8. Unannounced to participants, the effort contingencies periodically reversed. Reversals occurred after the response more likely to produce “easy” effort was chosen a cumulative number of times which was randomly selected to be between 5 and 9 for each reversal. Participants were instructed that their responses would affect the effort requirements in some way but were not informed of the specific nature of the task. Participants were not instructed to respond in any particular way other than to sample both choices.

**Figure 1.**
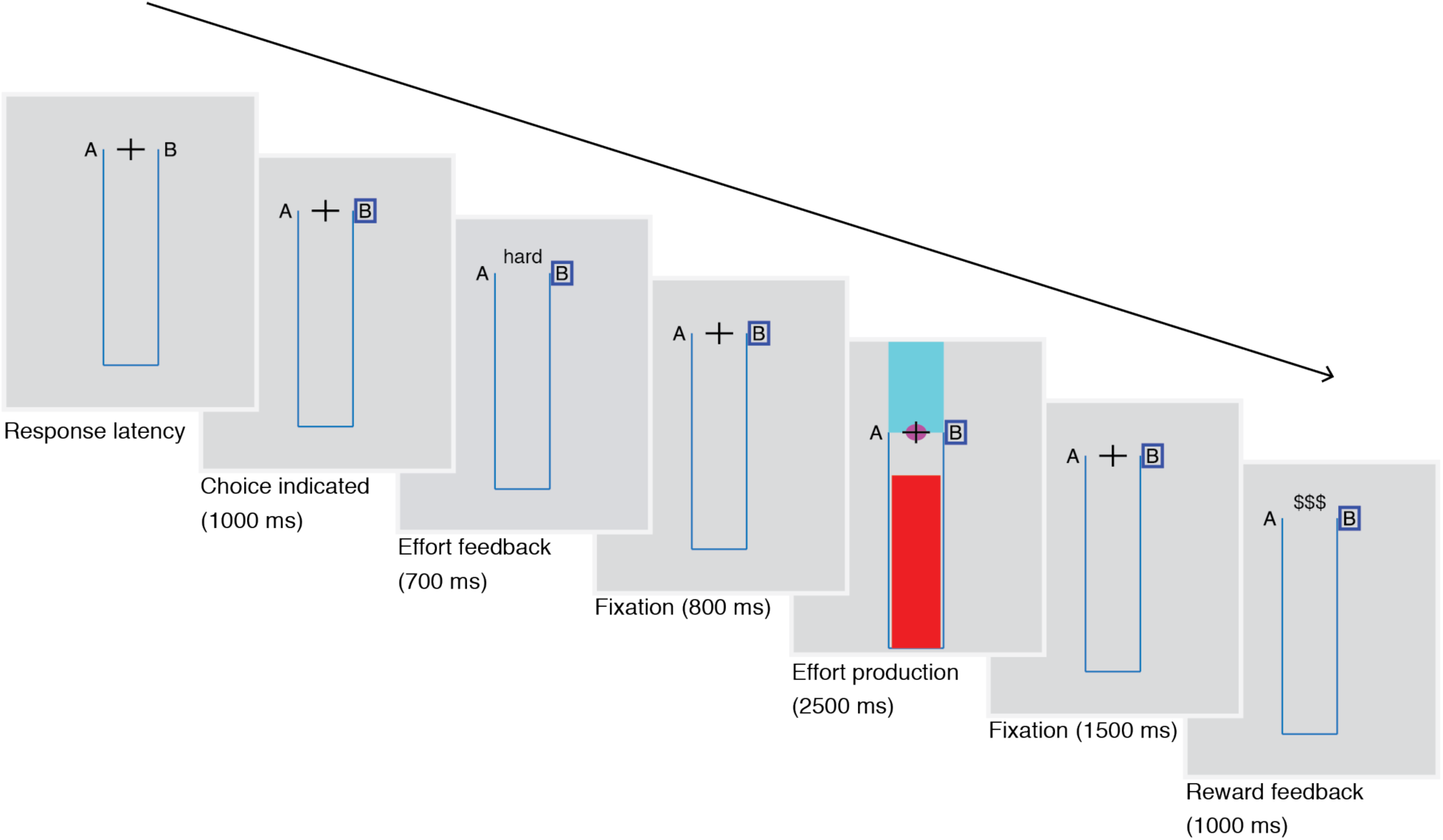
Stimuli. Participants initiated each trial by indicating a binary decision through button press. A box immediately appeared around either the letter A or B, corresponding to the choice options. After 1000 ms, feedback appeared to inform participants that their choice resulted in either “easy” or “hard” physical effort requirements for the upcoming EMG production task. A purple circle appeared over the target cross to cue the onset of the EMG production phase, during which participants performed isometric knee extension and the fluid level of a thermometer indicated quadriceps muscle activation. The circle shrank continuously, disappearing in 2.5 s to cue the end of the EMG production phase. Participants attempted to bring the fluid level above the target, represented by a cross, while remaining as close as possible to the target. However, a mask drawn above the target prevented participants from seeing the extent of errors that they made in overshooting the target. Instead, binary reinforcement feedback was provided 1.5 s after the EMG production phase ended, indicating whether or not participants had successfully exceeded the target while remaining sufficiently close.

After the effort feedback was removed from the display, the target cross reappeared for 800 ms. Subsequently, the effort production phase of the trial began. During this phase, the fluid level of the thermometer was drawn continuously to provide EMG feedback (see “Visual Feedback of EMG”). The fluid level increased with increasing EMG signal but represented the maximum signal for the trial, and thus never decreased. A purple circle was drawn under the target to cue the beginning of the effort production phase, and participants were instructed to keep their legs relaxed until they saw this cue. The circle shrank continuously during the course of the trial, disappearing in 2500 ms to signal the end of the effort production phase, at which point EMG feedback disappeared. Participants were instructed that in order to complete the task successfully, the final fluid level must exceed the target represented by the center of the cross. The target corresponded to 15% and 85% of MVC in the “easy” effort and “hard” effort conditions, respectively. Furthermore, participants were instructed to keep the fluid level as close as possible to the target, thus their goal was to always overshoot the target but to minimize the extent of overshoot. Participants were instructed to relax their legs as soon as possible after reaching the target, as the fluid level did not decrease during a trial. EMG feedback was withheld above the target by a mask drawn on the top of the thermometer. This prevented participants from seeing the extent of their overshoot errors, and performance feedback was instead provided by binary reinforcement.

At the end of the effort production phase, the EMG feedback and the mask disappeared. After 1500 ms of fixation, the target cross was replaced with either “$$$” or “XXX” to indicate a rewarded or failed trial, with a reward being indicated if the fluid level exceeded the target while remaining sufficiently close to it. Participants were instructed that they could earn up to an additional 10 CAD throughout the task according to the number of trials in which they received feedback indicating success. The error threshold for overshoot was adjusted using a 1-up-1-down adaptive staircase separately for the two effort conditions to ensure a 50% reinforcement rate overall for both conditions.

#### Control Condition

Each block began and ended with 12 control trials, during which the task was the same as the experimental condition except no reinforcement feedback was provided and the effort condition was deterministic and independent of participants’ responses. Both runs of 12 control trials consisted of 6 “easy” effort trials and 6 “hard” effort trials, with the trials of each effort condition occurring consecutively. The text “easy effort” or “hard effort” was displayed at the top of the screen continuously to cue the effort condition for all control trials. Participants were instructed to make a button press to initiate each trial, but that the choice was arbitrary and that the effort condition would always correspond to the cue at the top of the screen. In the first 12 control trials of each block, there was no mask drawn on the top of the thermometer, so participants could see their overshoot errors in order to practice the task more effectively. In the final 12 control trials of each block, the mask was drawn for each trial as in the experimental condition. The orders of “easy” and “hard” condition runs during the control trials were randomized and balanced across the four blocks for each participant.

#### Practice trials

Participants first performed a practice block to learn how to control the EMG feedback. As in the control trials, no reinforcement feedback was provided and the effort condition was cued to participants before each trial and independent of participants’ responses. The practice block began with 7 “easy” effort trials followed by 7 “hard” effort trials without the mask drawn at the top of the thermometer. Participants then performed 7 “easy” effort trials followed by 7 “hard” effort trials with the mask.

#### Behavioral Analysis

The effect of effort and reinforcement outcomes on behavioral choice was analyzed using logistic regression performed with the *Glmnet* package in R. The dependent variable was whether the participants’ choice on trial *n* corresponded to staying or switching from the choice on trial *n-1*, coded as 0 or 1. The independent variables were determined by the effort and reinforcement outcomes on trial *n-1*:

|  |  |
| --- | --- |
| *Effort:* | -1 for “easy” effort, 1 for “hard” effort |
| *Reward:* | -1 for non-reward, 1 for reward |
| *Effort/Reward interaction:* | *Effort\*Reward* |
| *Switch:* | 1 for all trials |

Logistic regression was calculated separately for each participant. Regularization was applied using L2-norm penalty. The penalty constant, *λ*, selected using leave-out cross validation. A value of 0.04297 was chosen as it is the largest value which minimizes the cross-validated misclassification error, averaged across subjects. The coefficients for *Effort, Reward*, and the interaction term were each submitted to 1-sample t-tests against zero.

### EEG Data Denoising

EEG data were preprocessed using the EEGLAB toolbox (see (Delorme and Makeig 2004a) for details), except for filtering which was performed using the MATLAB *filtfilt* function. Data, initially referenced to linked mastoids, were bandpass filtered using a second order Butterworth filter with a passband of 0.1–45 Hz. Channels with poor recording quality or excessive artifacts were identified with visual inspection and interpolated using spherical interpolation. EEG data were then re-referenced to the average scalp potential and interpolated electrodes were subsequently removed from the data before independent components analysis. Two epochs were extracted for each trial corresponding to effort condition feedback following the button press response and reinforcement feedback following the effort production phase. Continuous data were segmented into 2.5 second epochs time-locked to stimulus onset at 0 ms (time range: −1000 to +1500 ms). Data epochs containing artifacts other than blinks were removed by visual inspection. Subsequently, extended infomax independent component analysis was performed on each participant’s data (Delorme and Makeig 2004b). Components reflecting eye movements and blink artifacts were identified by visual inspection and subtracted by projection of the remaining components back to the voltage time series.

### Event Related Potential Analysis

#### Preprocessing and Trial averaging

We computed event related potentials (ERPs) on an individual participant basis by trial averaging EEG time series epochs after artifact removal. We selected trials corresponding to various feedback conditions in each task. For ERPs time locked to reinforcement feedback, we computed ERPs corresponding to “easy non-reward” (45.4 ± 6.9 trials) “easy reward” (46.3 ± 6.2 trials), “hard non-reward” (37.9 ± 8.4 trials), “hard reward” (45.6 ± 7.7 trials). In the control condition, participants performed the effort production task but did not receive any reinforcement feedback. We computed ERPs for the “control easy” (41.5 ± 5.9 trials) and “control hard” (38.5 ± 5.5 trials) conditions time locked to the moment where reinforcement feedback would have been delivered in the experimental condition. For ERPs corresponding to reinforcement feedback and the control condition, we excluded all trials in which the visual EMG feedback did not reach the target, as in this case a non-reward outcome was evident before the reinforcement feedback was delivered. We also extracted ERPs time locked to the effort condition feedback, which indicated the upcoming effort requirements after each button press but before the participant performed the EMG production task (“easy feedback” 94.9 ± 10.2 trials and “hard feedback” 92.4 ± 13.5 trials). All ERPs were baseline corrected by subtracting the average voltage in the 100 ms period immediately prior to stimulus onset. Finally, ERPs were low-pass filtered with a cutoff frequency of 30 Hz.

#### ROI selection

Because it is well established that reinforcement feedback elicits a feedback related negativity (FRN), and our hypothesis concerns the effect of physical effort on the FRN, we selected our electrode and time window of interest to maximize the contrast between the ERP responses to reward and non-reward feedback, irrespective of effort condition. For each subject, we computed the difference wave between ERPs elicited by reward feedback and non-reward feedback (reward minus non-reward), and selected our ROI using the grand average difference waves across subjects. We found that the largest peak occurred at electrode FCz at 234 ms after feedback onset. Thus we selected electrode FCz to analyze the feedback related negativity, which is consistent with previous work including our own (Miltner, Braun, and Coles 1997; Holroyd and Krigolson 2007; Pfabigan et al. 2011; Palidis, Cashaback, and Gribble 2019). We took the average ERP amplitude within a 50 ms window centered around 234 ms as a measure of the FRN.

#### Statistical Analysis

We analyzed the feedback related negativity using a temporal ROI approach by submitting the average ERP amplitude from 209–259 ms after feedback onset to statistical testing. For the neural response to reinforcement feedback, we performed a 2×2 repeated measures ANOVA using the MATLAB *ranova* function. The factors were reward outcome (levels: non-reward, reward) and effort condition (levels: easy, hard). Our time window was not defined a priori but rather selected to maximize the difference between reward and non-reward.

Without accounting for multiple comparisons across all possible time windows, the main effect of reward outcome is biased and the significance is inflated. However, our hypotheses concern the effect of effort and the interaction between effort and reward, with respect to which our ROI selection is blind and therefore unbiased.

We tested for artefacts related to the isometric leg extension using the difference wave computed between the “easy control” and “hard control” ERPs. We submitted the mean value of the difference wave between 209–259 ms to a one sample t-test against zero. These ERPs were aligned to the moment when reinforcement feedback would have been delivered in the experimental condition, but instead the target cross simply disappeared briefly. Participants were told that they would not receive feedback in this condition and thus did not expect a possible reward.

We tested for an FRN elicited by effort condition feedback using the difference wave computed between “easy feedback” and “hard feedback” ERPs. These ERPs were aligned to feedback indicating effort condition on each trial, occurring after the button press response but before the EMG production phase. We submitted the mean value of the difference wave between 209–259 ms to a one sample t-test against zero.

Because we observed effects outside of the time period of the FRN, we also performed statistical tests without averaging within a temporal ROI, instead testing each sample between 100-600ms after feedback onset. We selected this time window as it is wide enough to capture effects outside of the of the FRN, yet constrained to a range during which ERPs are likely to be affected by feedback processing (Glazer et al. 2018). We corrected significance values for multiple comparisons across time using the Benjamini-Hochberg procedure for estimating the false discovery rate (FDR), implemented by the MATLAB *mafdr* function. To analyze the neural response to reinforcement feedback, we used 2×2 repeated measures ANOVA with factors reward outcome and effort condition. We used one sample t-tests against zero on the difference waves computed between “easy feedback” and “hard feedback” ERPs, aligned to feedback indicating effort condition after each button press. We also used one sample t-tests against zero on the difference waves computed between “easy control” and “hard control” ERPs, aligned to the moment when reinforcement feedback would have been delivered.

#### Scalp Distributions

Scalp distributions were plotted with the EEGLAB *topoplot* function using the mean amplitude of difference waves within specified time windows, averaged across subjects.

## Results

### Behavioral Results

Participants made binary decisions which probabilistically determined the effort requirements for each trial. Participants underwent the “hard effort” condition in 49.6% (std: 4.8%) of trials. Reward was delivered if EMG feedback exceeded a target level while staying sufficiently close to the target. Participants received reward in 49.4% (std: 0.02%) of trials. We performed logistic regression for each subject to predict switching of responses between trials *n-1* and *n*, with the effort condition and reward outcome on trial *n-1* as the predictors. Figure 2b shows the coefficients estimated for each subject, and Figure 2a shows the proportion of trials after which participants switched responses for the different reward and effort outcomes. We found that the coefficients for the effect of effort on switching were significantly greater than zero (one-sample *t* test; t(17) = 2.263, p = 0.037). The coefficients for the effect of reward were not reliably different from zero (t(17) = −0.871, p = 0.3959), nor were the coefficients for the interaction term (t(17) = 0.252, p = 0.8043).

**Figure 2.**
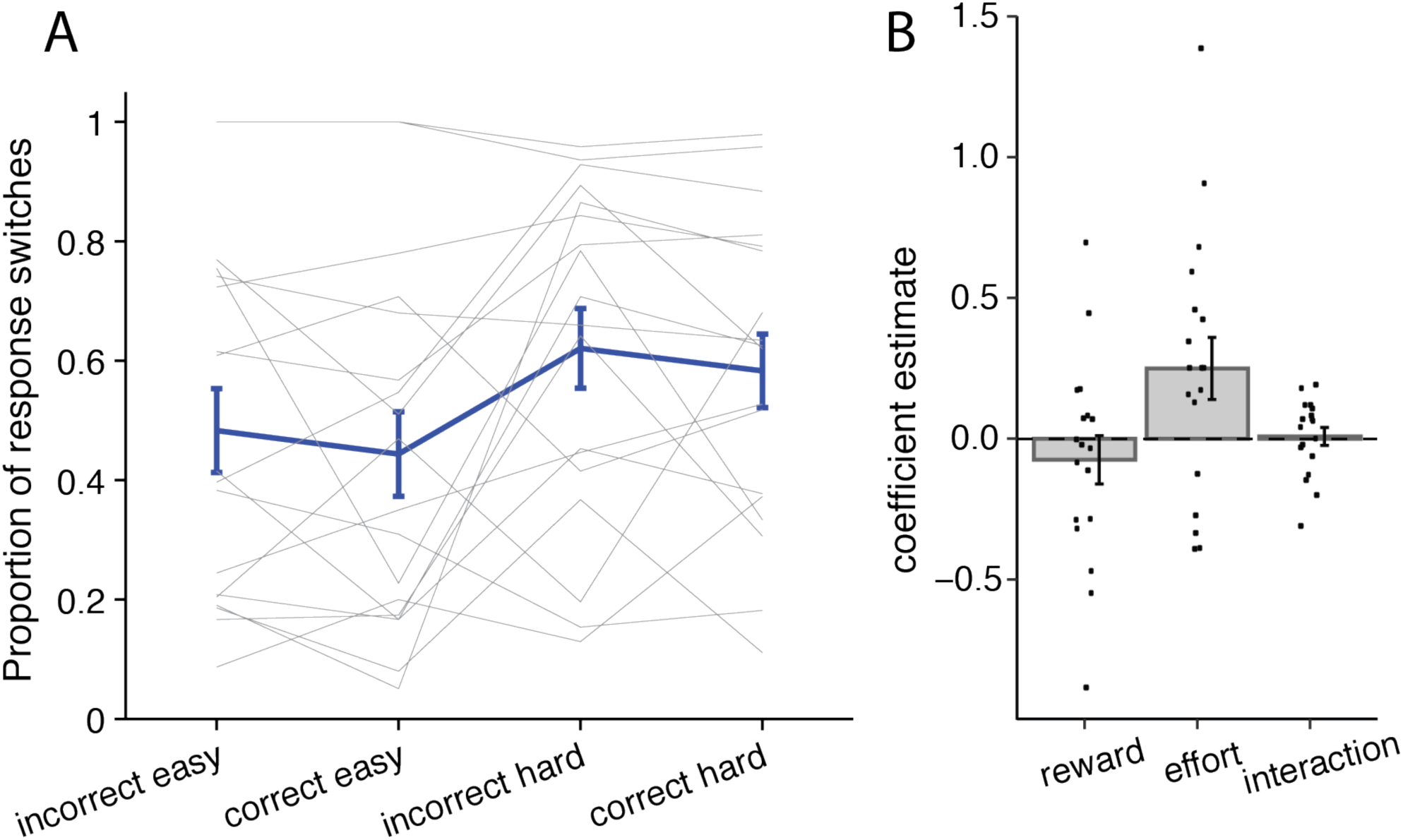
Participants switch responses more frequently after hard physical effort than easy effort. **A**, The proportion of trials on which participants switched responses between trial *n-1* and trial *n* for the different reward outcomes and effort conditions on trial *n-1* (Error bars: ± SEM). **B**, Coefficients estimated using logistic regression to predict response switching for each participant (Bars indicate mean: ± 1 SEM). Predictors were reward outcome, effort condition, reward/effort interaction. The effort term was significantly greater than zero (p = .037), indicating that participants were more likely to switch responses after hard effort than easy effort.

### ERP Results

#### Feedback Related Negativity Time Window

Figure 3a shows the ERPs elicited by reinforcement feedback, while Figure 3b shows the effects of effort and reward on ERP amplitude during the FRN time window. We analyzed the neural response to reinforcement feedback by performing 2×2 repeated measures ANOVA on the average ERP amplitude within our FRN time window. We found that voltage was lower in the “hard effort” condition compared to the “easy effort” condition (main effect, F(1,17) = 12.41, p = 0.0026). We also found that voltage was more positive for reward compared to non-reward (main effect, F(1,17) = 49.8, p < 0.001). This effect is inflated as we selected our time window of interest to maximize the effect of reward. However, the effect is still reliable after Bonferroni correction for all possible placements of a 50 ms window within the 1.5 second period of our epoch after feedback presentation (corrected p = 0.0014). We found no reliable interaction between reward and effort (F(1,17) = 1.09, p = 0.31). In the control condition, participants performed the EMG production task but the effort condition was pre-determined and no reinforcement feedback was provided. We aligned neural responses to the moment in the trial when reinforcement feedback would have been provided in the experimental condition. We found that the “control easy” - “control hard” difference wave was not reliably different from zero during the FRN time window (mean difference: 0.275 µv, SD: 1.41, t(17) = 0.826, p = 0.420).

**Figure 3.**
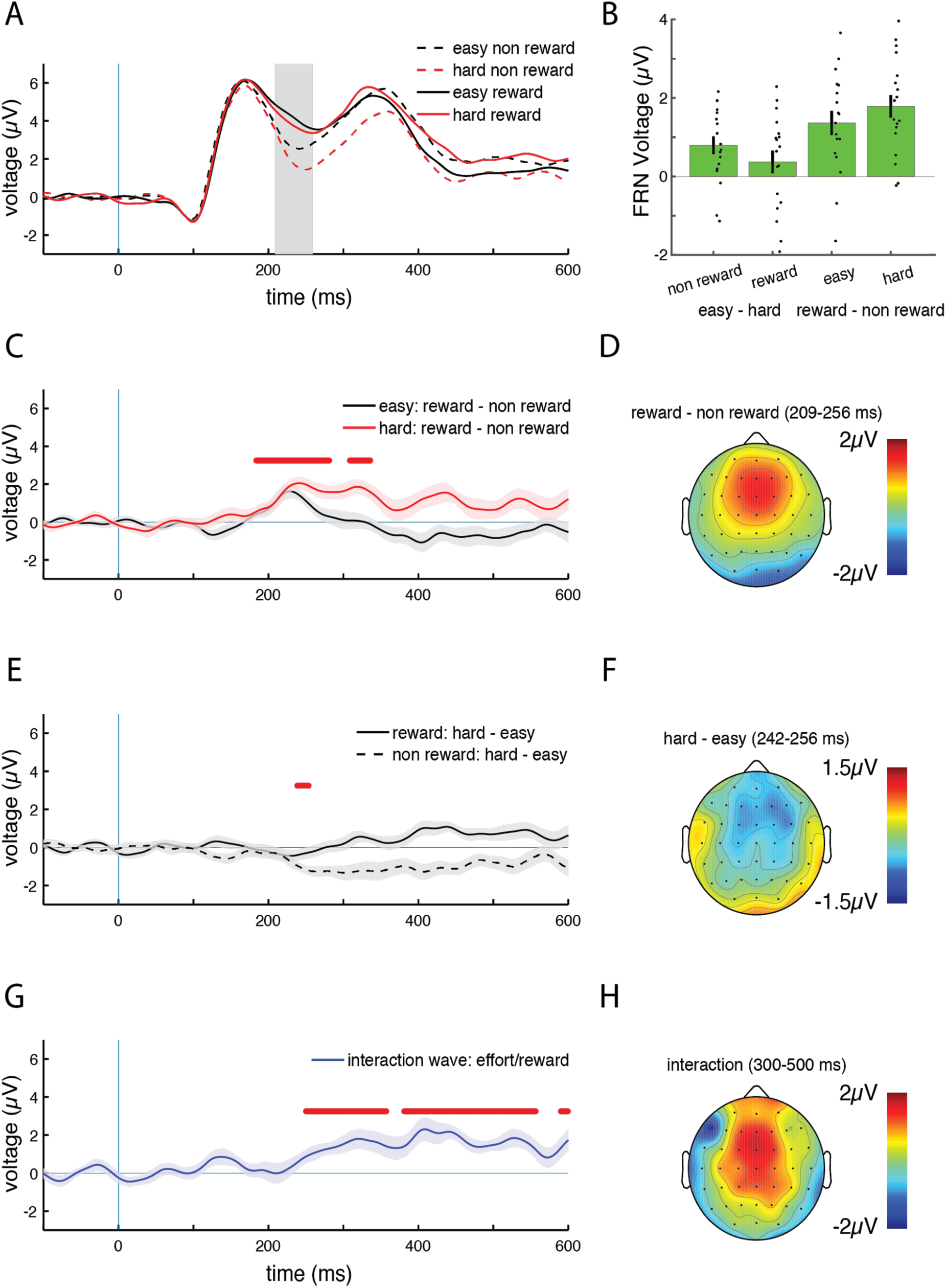
ERPs elicited by reinforcement feedback. **A**, *Trial averaged ERPs recorded from electrode FCz aligned to reinforcement feedback presentation (0 ms: vertical blue line), selected for reinforcement outcome (reward or non reward) and the physical effort requirement on that trial (“easy” or “hard”). Shaded area represents time window for feedback related negativity (frn) analysis (209-259 ms).****B***, *Effects of effort and reward on average ERP voltage within the FRN time window. The effect of effort (“easy” ERP amplitude - “hard” ERP amplitude) is shown separately for non reward and reward outcomes. The effect of reward (reward ERP amplitude - non reward ERP amplitude) is shown separately for “easy” and “hard” effort conditions.* (Error bars: ± SEM) **C**, Mean difference waves computed as reward ERP - non reward ERP, separately for the “easy” and “hard” effort conditions (Shaded region: ± SEM). Red markers indicate time points between 100-600 ms with significant main effect of reward outcome (p < 0.05, FDR corrected). **D**, Scalp distribution of reward - non reward ERPs, irrespective of effort condition, between 209-256 ms. **E**, Mean difference waves computed as “hard” effort ERP - “easy” effort ERP, separately for the reward and non reward effort conditions (Shaded region: ± SEM). Red markers indicate time points between 100-600 ms with significant main effect of effort condition (p < 0.05, FDR corrected). **F**, Scalp distribution of “hard” effort - “easy” effort ERPs, irrespective of reward outcome, between 209-256 ms. **G**, Mean interaction wave computed as (“hard” reward ERP - “hard” non reward ERP) - (“easy” reward ERP - “easy” non reward ERP). Shaded region: ± SEM. Red markers indicate time points between 100-600 ms with significant interaction effect of reward outcome (p < 0.05, FDR corrected). **H**, Scalp distribution of interaction wave between 300-500 ms.

After participants produced binary decisions by button press, feedback was provided to indicate the resulting effort condition for the current trial. Figure 4a shows the ERPs elicited on each trial by feedback indicating the effort condition. We found that the “easy feedback” - “hard feedback” difference wave was not reliably different from zero during the FRN time window (mean difference: −0.277 µv, SD: 0.953, t(17) = −1.2317, p = 0.235).

**Figure 4.**
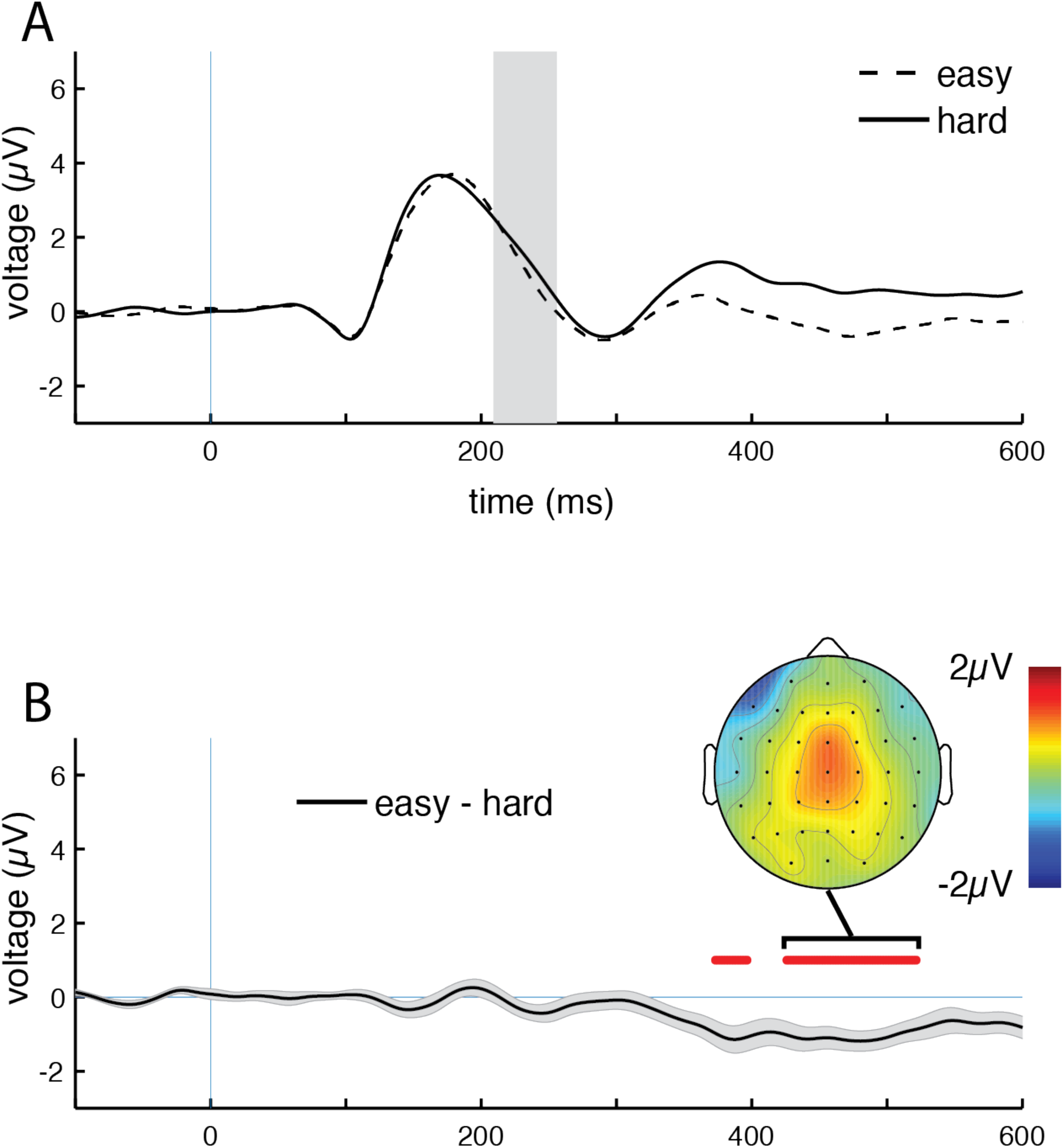
ERPs elicited by effort condition feedback. **A**, *Trial averaged ERPs recorded from electrode FCz aligned to effort condition feedback presentation (0 ms: vertical blue line), selected for physical effort condition (“easy” or “hard”). Shaded area represents time window for feedback related negativity (frn) analysis (209-259 ms).* ***B***, Mean difference waves computed as “easy feedback” ERP - “hard feedback” ERP (Shaded region: ± SEM). Red markers indicate time points between 100-600 ms significantly different from zero (p < 0.05, FDR corrected). Inset: Scalp distribution of difference wave between 426-521 ms.

#### Sample-wise analysis

We performed the same statistical tests used for the FRN time window, but for each individual time point 100–600 ms post feedback onset. P-values are corrected for multiple comparisons across time points using FDR. In response to reinforcement feedback, we found reliable main effects of reward outcome between 184–336 ms after feedback onset (Figure 3c, ranges for significant time points: F = [7.99 52.99], p = [.0001 .045], uncorrected p = [<0.0001 .012]). We found main effects of effort condition 238–254 ms after feedback onset (Figure 3e, ranges for significant time points: F = [13.74 14.45], p = [0.050 0.050], uncorrected p = [0.0011 .0018]). We found effort/reward interaction effects starting 250 ms after feedback onset and up to 600 ms, the end of our time window for statistical testing (Figure 3g, ranges for significant time points: F = [5.73 38.94], p = [0.0008 0.048], uncorrected p = [<0.0001 0.028]). We found no reliable differences between the “control easy” and “control hard” ERPs (ranges for all timepoints between 100-600 ms: t = [-1.56 1.42], uncorrected p = [.139 1.00]. We found significant differences between the “easy feedback” and “hard feedback” ERPs between 373-522 ms after effort feedback (Figure 4b, ranges for significant time points: t = [−4.31 −2.81], p = [0.015 0.049], uncorrected p = [0.0005 0.012]). Although we only performed statistical testing up to 600 ms to avoid sacrificing statistical power, we observed that the “easy feedback” - “hard feedback” difference wave remains at least 1 standard error below zero until 1018 ms post-feedback.

## Discussion

Participants were more likely to switch responses after choices that led to “hard” effort than “easy” effort, suggesting that they adapted behavior to reduce physical effort in response to uncertain outcomes. At the end of each trial, binary reinforcement feedback indicated whether participants achieved a monetary reward which depended on precisely producing a target level of EMG activity. Unsurprisingly, reinforcement feedback elicited a robust feedback related negativity (FRN) response. The FRN was characterized by a negativity over the frontocentral scalp in response to feedback indicating non-reward relative to a positivity occurring in response to reward. The effect of preceding effort during the FRN window was opposite that of reward, with reinforcement outcomes which required more effort producing a lower ERP amplitude. This finding is consistent with the hypothesis that the FRN encodes a prediction error which is computed not only using the value of the reward outcome but rather based on a measure of subjective utility, which is a function of reward outcome discounted by the physical effort required to obtain that outcome.

Within the temporal window containing the FRN, we found oppositional main effects of reward and effort on the amplitude of the neural response. This temporal ROI approach allowed us to test for effects of effort with high statistical power. However, a sample-wise analysis revealed more complicated dynamics, with reward and effort producing main effects and an interaction that only partly overlapped in time. After reinforcement feedback was delivered, an effect of reward outcome first emerged with a latency of 184 ms, which remained significant while an additional main effect of preceding effort emerged at 238 ms. Finally, a sustained interaction between reward and effort first occurred around 250 ms after feedback onset. This onset of the interaction is still within the typical range of the FRN, but the effect persisted upwards of 600 ms, beyond the time range typically associated with the FRN. All of these effects had a similar frontocentral scalp distribution. These dynamics suggest a process whereby upon receiving reward feedback, the medial frontal cortex first encodes the immediate reward outcome and subsequently integrates signals related to the preceding effort. This process culminates in an interaction whereby the effect of reward outcome depends on the preceding effort. While the initial effect of reward outcome and the later interaction are clear, the main effect of effort occurring during the FRN prior to the interaction effect is much weaker. It is possible that this main effect of effort is driven primarily by the emerging interaction effect, and that there is simply not enough statistical power to detect the interaction during that time interval.

The neural interaction between effort and reward was characterized by a larger, more sustained effect of reinforcement outcome when the preceding effort was high. Analogous interactions have been observed behaviorally in animals and humans, in which the same reward produces stronger reinforcement when it requires more effort to obtain (Inzlicht, Shenhav, and Olivola 2018; Lydall, Gilmour, and Dwyer 2010; Zentall 2010; Clement et al. 2000). Unfortunately, we were not able to assess such a behavioral interaction in the current study as there was no effect of reward outcome on the binary decisions that participants made on each trial. This was not surprising as reward outcome was not determined by these decisions but rather by performance on the EMG production task. The binary decisions only determined the effort required, and the reinforcement threshold was controlled to produce approximately equal reward rate in both effort conditions.

After participants produced binary responses, feedback indicated the resulting physical effort condition for the subsequent EMG production portion of the trial. In line with theoretical accounts of the FRN as a temporal difference learning prediction error signal, stimuli which predict aversive outcomes or economic loss typically elicit FRN responses (Mulligan and Hajcak 2018). Thus, we predicted that effort feedback might elicit an FRN component as a learning signal for effort minimization. However, we observed no FRN component when comparing the ERP responses to feedback indicating “easy” or “hard” effort trials. Rather, effort modulated the FRN in response to reinforcement feedback at the end of the trial. This suggests that physical effort is not immediately treated by the reinforcement learning system as a loss or a punishing stimulus. Rather, effort information can be maintained during the course of an action and incorporated with reward information at the time of outcome evaluation. We often undertake protracted tasks for which the effort requirements and ultimate payoffs are uncertain. It may not be efficient to punish the value representation of a task every time an unexpected effort is encountered, as the eventual payoff may be well worth the effort. Instead, it may be more efficient to integrate effort over the entire course of an undertaking and evaluate the cost and benefit simultaneously when the final outcome is observed. This process can also support interactions in which the effect of effort depends on reward which is only delivered later. Alternatively, some work suggests that we learn about effort requirements and reward separately and integrate them at the time of decision making (Skvortsova, Palminteri, and Pessiglione 2014; Hauser, Eldar, and Dolan 2017). It is likely that economic decision making and learning involves distributed hierarchical computations, and that it is possible to observe a distribution of signals with varying dependencies on effort, reward, and integrated utility throughout the brain (Hunt and Hayden 2017).

### Limitations

Participants adapted their behavior to reduce physical effort, but the behavioral effect of effort was variable and relatively weak. Participants were more likely to switch responses after choices that led to “hard” effort than “easy” effort. However, participants often switched responses after “easy” trials or stayed with responses that produced “hard” effort: on average, participants switched responses after 46.5% of “easy” trials and 60% of “hard” trials. Furthermore, negative coefficients for the effect of effort on switching were estimated for several participants. The relatively weak and highly variable effects of effort is consistent with the notion that although effort is generally treated as a cost which is minimized, in many cases people are undeterred by effort or even purposefully select more effortful options (Inzlicht et al. 2018; Eisenberger 1992). These effects are often attributed to state-dependent learning in which reinforcement outcomes are evaluated relative to the value of the current state. In the current study, variable reinforcement outcomes were only evaluated after effort production, and thus may have been more valuable when received after a costly, high effort action. Other details of the task may have affected effort related choice. Unlike some previous studies of effort minimization, participants were not instructed to avoid effort. Furthermore, success in the task was not dependent on exerting effort which exceeded an unknown criterion. These features may enhance effort minimization but they could also conflate effort prediction errors with errors relative to the goals of the task at hand, which are also strongly represented in the ACC (Krigolson and Holroyd 2007; Fu et al. 2018; Ullsperger et al. 2014; Swick and Turken 2002).

Although the excellent temporal resolution offered by EEG proved instrumental in uncovering the dynamics of effort and reward processing in the brain, it invariably measures a mixture of signals from neurons with different response properties. Kennerley et al., 2011 identified diverse tuning to economic value across ACC, orbitofrontal cortex, and lateral prefrontal cortex, such that many neurons that are selective to value with opposite tunings will cancel out at the population level measured by EEG of FMRI. One notable exception was a subpopulation of ACC neurons which encoded value with positive tuning multiplexed across reward, effort, and delay variables. Interestingly, these neurons also tended to encode positive reward prediction errors. Furthermore, EEG measured at the scalp is difficult to localize and can represent mixtures of activity from entirely separate brain regions. Although the FRN is a well characterized response and convergent lines of evidence suggest a source in the ACC, we also report effects outside the typical time range of the FRN. These effects exhibit a medial-frontal scalp distribution which is similar to the FRN, however the neural source cannot be determined with certainty.

## References

Amiez, Céline, Jean-Paul Joseph, and Emmanuel Procyk. 2005. “Anterior Cingulate Error-Related Activity Is Modulated by Predicted Reward.” The European Journal of Neuroscience 21 (12): 3447–52.

Becker, Michael P. I., Alexander M. Nitsch, Wolfgang H. R. Miltner, and Thomas Straube. 2014. “A Single-Trial Estimation of the Feedback-Related Negativity and Its Relation to BOLD Responses in a Time-Estimation Task.” The Journal of Neuroscience: The Official Journal of the Society for Neuroscience 34 (8): 3005–12.

Clement, Tricia S., Joann R. Feltus, Daren H. Kaiser, and Thomas R. Zentall. 2000. “‘work Ethic’ in Pigeons: Reward Value Is Directly Related to the Effort or Time Required to Obtain the Reward.” Psychonomic Bulletin & Review 7 (1): 100–106.

Croxson, Paula L., Mark E. Walton, Jill X. O’Reilly, Timothy E. J. Behrens, and Matthew F. S. Rushworth. 2009. “Effort-Based Cost-Benefit Valuation and the Human Brain.” The Journal of Neuroscience: The Official Journal of the Society for Neuroscience 29 (14): 4531–41.

Delorme, Arnaud, and Scott Makeig. 2004a. “EEGLAB: An Open Source Toolbox for Analysis of Single-Trial EEG Dynamics Including Independent Component Analysis.” Journal of Neuroscience Methods 134 (1): 9–21.

Delorme, Arnaud. 2004b. “EEGLAB: An Open Source Toolbox for Analysis of Single-Trial EEG Dynamics Including Independent Component Analysis.” Journal of Neuroscience Methods 134 (1): 9–21.

Denk, F., M. E. Walton, K. A. Jennings, T. Sharp, M. F. S. Rushworth, and D. M. Bannerman. 2005. “Differential Involvement of Serotonin and Dopamine Systems in Cost-Benefit Decisions about Delay or Effort.” Psychopharmacology 179 (3): 587–96.

Glazer, James E., Nicholas J. Kelley, Narun Pornpattananangkul, Vijay A. Mittal, and Robin Nusslock. 2018. “Beyond the FRN: Broadening the Time-Course of EEG and ERP Components Implicated in Reward Processing.” International Journal of Psychophysiology: Official Journal of the International Organization of Psychophysiology 132 (Pt B): 184–202.

Graybiel, Ann M. 2008. “Habits, Rituals, and the Evaluative Brain.” Annual Review of Neuroscience 31: 359–87.

Hauser, Tobias U., Eran Eldar, and Raymond J. Dolan. 2017. “Separate Mesocortical and Mesolimbic Pathways Encode Effort and Reward Learning Signals.” Proceedings of the National Academy of Sciences of the United States of America 114 (35): E7395–7404.

Holroyd, Clay B., and G. H. 2002. “The Neural Basis of Human Error Processing: Reinforcement Learning, Dopamine, and the Error-Related Negativity.” Psychological Review 109 (4): 679–709.

Holroyd, Clay B., and Olave E. Krigolson. 2007. “Reward Prediction Error Signals Associated with a Modified Time Estimation Task.” Psychophysiology 44 (6): 913–17.

Holroyd, Clay B., Olav E. Krigolson, and Seung Lee. 2011. “Reward Positivity Elicited by Predictive Cues.” Neuroreport 22 (5): 249–52.

Hosking, Jay G., Stan B. Floresco, and Catharine A. Winstanley. 2015. “Dopamine Antagonism Decreases Willingness to Expend Physical, but Not Cognitive, Effort: A Comparison of Two Rodent Cost/benefit Decision-Making Tasks.” Neuropsychopharmacology: Official Publication of the American College of Neuropsychopharmacology 40 (4): 1005–15.

Hunt, Laurence T., and Benjamin Y. Hayden. 2017. “A Distributed, Hierarchical and Recurrent Framework for Reward-Based Choice.” Nature Reviews. Neuroscience 18 (3): 172–82.

Inzlicht, Michael, Amitai Shenhav, and Christopher Y. Olivola. 2018. “The Effort Paradox: Effort Is Both Costly and Valued.” Trends in Cognitive Sciences 22 (4): 337–49.

Ito, Shigehiko, Veit Stuphorn, Joshua W. Brown, and Jeffrey D. Schall. 2003. “Performance Monitoring by the Anterior Cingulate Cortex during Saccade Countermanding.” Science 302 (5642): 120–22.

Kennerley, Steven W., Timothy E. J. Behrens, and Jonathan D. Wallis. 2011. “Double Dissociation of Value Computations in Orbitofrontal and Anterior Cingulate Neurons.” Nature Neuroscience 14 (12): 1581–89.

Klein-Flügge, Miriam C., Steven W. Kennerley, Karl Friston, and Sven Bestmann. 2016. “Neural Signatures of Value Comparison in Human Cingulate Cortex during Decisions Requiring an Effort-Reward Trade-Off.” The Journal of Neuroscience: The Official Journal of the Society for Neuroscience 36 (39): 10002–15.

Kurniawan, Irma Triasih, Marc Guitart-Masip, and Ray J. Dolan. 2011. “Dopamine and Effort-Based Decision Making.” Frontiers in Neuroscience 5 (June): 81.

Kurniawan, I. T., B. Seymour, D. Talmi, W. Yoshida, N. Chater, and R. J. Dolan. 2010. “Choosing to Make an Effort: The Role of Striatum in Signaling Physical Effort of a Chosen Action.” Journal of Neurophysiology 104 (1): 313–21.

Lydall, Emma S., Gary Gilmour, and Dominic M. Dwyer. 2010. “Rats Place Greater Value on Rewards Produced by High Effort: An Animal Analogue of the ‘effort Justification’ Effect.” Journal of Experimental Social Psychology 46 (6): 1134–37.

Miltner, W. H., C. H. Braun, and M. G. Coles. 1997. “Event-Related Brain Potentials Following Incorrect Feedback in a Time-Estimation Task: Evidence for a ‘Generic’ Neural System for Error Detection.” Journal of Cognitive Neuroscience 9 (6): 788–98.

Morel, Pierre, Philipp Ulbrich, and Alexander Gail. 2017. “What Makes a Reach Movement Effortful? Physical Effort Discounting Supports Common Minimization Principles in Decision Making and Motor Control.” PLoS Biology 15 (6): e2001323.

Mulligan, Elizabeth M., and Greg Hajcak. 2018. “The Electrocortical Response to Rewarding and Aversive Feedback: The Reward Positivity Does Not Reflect Salience in Simple Gambling Tasks.” International Journal of Psychophysiology: Official Journal of the International Organization of Psychophysiology 132 (Pt B): 262–67.

Palidis, Dimitrios J., Joshua G. A. Cashaback, and Paul L. Gribble. 2019. “Neural Signatures of Reward and Sensory Error Feedback Processing in Motor Learning.” Journal of Neurophysiology 121 (4): 1561–74.

Pfabigan, Daniela M., Johanna Alexopoulos, Herbert Bauer, and Uta Sailer. 2011. “Manipulation of Feedback Expectancy and Valence Induces Negative and Positive Reward Prediction Error Signals Manifest in Event-Related Brain Potentials.” Psychophysiology 48 (5): 656–64.

Porter, Blake S., Kristin L. Hillman, and David K. Bilkey. 2019. “Anterior Cingulate Cortex Encoding of Effortful Behavior.” Journal of Neurophysiology 121 (2): 701–14.

Puig, M. Victoria, and Earl K. Miller. 2012. “The Role of Prefrontal Dopamine D1 Receptors in the Neural Mechanisms of Associative Learning.” Neuron 74 (5): 874–86.

Rangel, Antonio, and Todd Hare. 2010. “Neural Computations Associated with Goal-Directed Choice.” Current Opinion in Neurobiology 20 (2): 262–70.

Schultz, Wolfram. 2006. “Behavioral Theories and the Neurophysiology of Reward.” Annual Review of Psychology 57: 87–115.

Schweimer, Judith, Simone Saft, and Wolfgang Hauber. 2005. “Involvement of Catecholamine Neurotransmission in the Rat Anterior Cingulate in Effort-Related Decision Making.” Behavioral Neuroscience 119 (6): 1687–92.

Shadmehr, Reza, Helen J. Huang, and Alaa A. Ahmed. 2016. “A Representation of Effort in Decision-Making and Motor Control.” Current Biology: CB 26 (14): 1929–34.

Skvortsova, Vasilisa, Stefano Palminteri, and Mathias Pessiglione. 2014. “Learning to Minimize Efforts versus Maximizing Rewards: Computational Principles and Neural Correlates.” The Journal of Neuroscience: The Official Journal of the Society for Neuroscience 34 (47): 15621–30.

Vezoli, Julien, and Emmanuel Procyk. 2009. “Frontal Feedback-Related Potentials in Nonhuman Primates: Modulation during Learning and under Haloperidol.” The Journal of Neuroscience: The Official Journal of the Society for Neuroscience 29 (50): 15675–83.

Walsh, Matthew M., and John R. Anderson. 2012a. “Learning from Experience: Event-Related Potential Correlates of Reward Processing, Neural Adaptation, and Behavioral Choice.” Neuroscience and Biobehavioral Reviews 36 (8): 1870–84.

Walsh, Matthew M. 2012b. “Learning from Experience: Event-Related Potential Correlates of Reward Processing, Neural Adaptation, and Behavioral Choice.” Neuroscience and Biobehavioral Reviews 36 (8): 1870–84.

Walton, Mark E., David M. Bannerman, Karin Alterescu, and Matthew F. S. Rushworth. 2003. “Functional Specialization within Medial Frontal Cortex of the Anterior Cingulate for Evaluating Effort-Related Decisions.” The Journal of Neuroscience: The Official Journal of the Society for Neuroscience 23 (16): 6475–79.

Walton, M. E., S. W. Kennerley, D. M. Bannerman, P. E. M. Phillips, and M. F. S. Rushworth. 2006. “Weighing up the Benefits of Work: Behavioral and Neural Analyses of Effort-Related Decision Making.” Neural Networks: The Official Journal of the International Neural Network Society 19 (8): 1302–14.

Williams, Ziv M., George Bush, Scott L. Rauch, G. Rees Cosgrove, and Emad N. Eskandar. 2004. “Human Anterior Cingulate Neurons and the Integration of Monetary Reward with Motor Responses.” Nature Neuroscience 7 (12): 1370–75.

Zentall, Thomas R. 2010. “Justification of Effort by Humans and Pigeons: Cognitive Dissonance or Contrast?” Current Directions in Psychological Science 19 (5): 296–300.

